# Insights into the mechanism of mycelium transformation of *Streptomyces toxytricini* into pellet

**DOI:** 10.1101/2023.02.01.526547

**Authors:** Punit Kumar, Khushboo, Deepanshi Rajput, Kashyap Kumar Dubey

## Abstract

Formation of the mycelial pellet in submerged cultivation of *Streptomycetes* is unwanted in industrial fermentation processes as it imposes mass transfer limitations, changes in the rheology of a medium, and affects the production of secondary metabolites. Though detailed information is not available about the factors involved in regulating mycelial morphology but it is studied that culture conditions and genetic information of strain play a key role. Moreover, the proteomic study has revealed the involvement of low molecular weight proteins such as; DivIVA, FilP, ParA, Scy, and SsgA proteins in apical growth and branching of hyphae which results in the establishment of the mycelial network. The present study proposes the mechanism of pellet formation of *Streptomyces toxytricini* (NRRL B-5426) with the help of microscopic and proteomic analysis. The microscopic analysis revealed that growing hyphae followed a certain organized path of growth and branching, which was further converted into the pellet, and proteomic analysis revealed the production of low molecular weight proteins, which possibly participate in the regulation of pellet morphology.

## 1. Introduction

During submerged fermentation conditions, filamentous microorganisms exhibit morphologies between diffused mycelia and pellets as per the culture conditions and type of microbial strain. It is evident that mycelial morphologies are associated with the production of the desired natural product. Moreover, the pellet morphologies affect the rheology of a medium, heterogeneous mass transfer, and downstream processing (Rioseras et al., 2014; Wang et al., 2017). Therefore, regulation of pellet morphology is necessary for industrial processes.

Studies conducted at the molecular level suggested the involvement of many genes and proteins that control the mycelium transformation or morphological differentiation of Streptomyces (Vassallo et al., 2020; Wu et al., 2020). The growth of *Streptomyces* takes place from spore into vegetative hyphae which is under the regulation of AMP receptor protein Crp (Derouaux et al., 2004). Further growth of the hyphae takes place by tip extension and branching, where polarity protein DivIVA plays a key role (Flardh et al., 2012). It was suggested that during the growth of hyphae, the development of cross-walls also takes place, which protects hyphae from fission and forms a multinucleated compartment (Jakimowicz and van Wezel, 2012). Moreover, many protein complexes of *Streptomyces* like Scy, Tat secretion system, SsgA, and CslA are also found to be associated with apical tip growth of *Streptomyces* (Hempel et al., 2012; Holmes et al., 2013; Noens et al., 2007; Willemse et al., 2012; Xu, 2008). It was reported that SsgA is involved in the identification of a location for the development of septum and germination site (Noens et al., 2007). Interestingly, Scy, ParA, and FilP proteins are assumed to interact with DivIVA to control the apical growth of mycelia (Ditkowski et al., 2013; Holmes et al., 2013). One more protein HyaS has been found conserved in streptomycetes and found to be associated with the regulation of pellet morphology by maintaining hyphal contacts (Koebsch et al., 2009).

Moreover, it was also suggested that some extracellular materials (proteins, sugars, and DNA) work as adhesives and are involved in the formation of pellet-like structures and provide protection. Extracellular DNA, hyaluronic acid, teichoic acids, and CslA have been found to be associated with the pellet morphology of *Streptomyces* (Kim and Kim, 2004; Ultee et al., 2020). It was found that cellulose synthase-like protein CslA was also identified near the hyphal tip which was involved in growth and structural transformation. It was also studied that it interacts with DivIVA and regulate the formation of biopolymer with glycosidic bonds near hyphal tips. The physical phenomenon, such as oxygen transfer and shear rate, also influence the morphology of *Streptomyces* (Ribeiro et al., 2021). Directive and quick-acting approaches, like the addition of microparticles and macroparticles, have also been used to regulate mycelial morphology (Böl et al., 2020; Yue et al., 2021).

*Streptomyces* genus is a group of Gram’s positive bacteria which are one of the most explored microorganisms at the commercial scale for the synthesis of natural products. During submerged fermentation, these microorganisms represent pellet morphology (Hobbs et al., 1989). *Streptomyces toxytricini* is recognized for the production of lipase-inhibitory natural product lipstatin; which is used in an FDA-approved anti-obesity drug (Kumar and Dubey, 2015,16). Like other species of the genus *Streptomyces toxytricini* also exhibits pellet morphology in submerged fermentation (Kumar and Dubey, 2017). In the previous report, authors have reported that culture conditions (pH of culture medium, medium composition, agitation rate, and inoculum size) influence the pellet size of *S. toxytricini* (Kumar and Dubey, 2017), but the mechanism of mycelial transformation into pellet has not been elaborated. Though pellet size was reduced in with change in culture conditions but biomass formation was also reduced. Thus in order to reduce the pellet size or to maintain dispersed mycelial morphology without affecting the biomass, mechanism of pellet formation must be studied. In the present study authors made an attempt to microscopic analysis of hyphae growth and branching which further transformed into the pellet. Further, partial analysis of proteins (produced by bacteria in pellet and culture medium) was performed to understand the role of proteins in pellet formation.

## 2. Materials and methods

### 2.1 Microorganism and growth conditions

*S. toxytricini* NRRL B-5426 strain was obtained from the Agricultural Research Service (NRRL), Department of Agriculture, United States. The bacterium was grown in a shake flask as per the conditions mentioned in (Kumar and Dubey, 2017). Incubation temperature was maintained at 27.5°C till the appearance of visible colonies. Well-grown colonies of *S. toxytricini* were pink in colour, elevated, circular in shape, and having a specific odour.

### 2.2 Analysis of pellet morphology

For pellet morphology analysis, an inoculum of *S. toxytricini* was prepared by transferring a loopful of bacteria (*S. toxytricini*) from a Petri plate into a shake flask, modified from (Kumar and Dubey, 2017).

After incubation, pellets were settled down and centrifuged at 5000 rpm for 10 min to remove the culture medium. Aggregated pellets were washed thrice by 0.1M sodium phosphate buffer of pH 7.2±0.1 and stored for further processing. The Gram’s staining and methylene blues staining of pellets were performed to analyze morphology and shape. Visualization was done in a compound microscope. For scanning electron microscopy (SEM) of pellets, primary fixation of pellets was performed in 2.5% glutaraldehyde and 2% paraformaldehyde in 0.1M sodium phosphate buffer at pH 7.2±0.1. Further dehydration, fixation and coating, and SEM analysis were done at Advanced Instrumentation Research Facility, Jawaharlal Nehru University, New Delhi (India). For biomass analysis, pellets were kept for drying at 50 °C in a hot air oven till the appearance of constant weight.

### 2.3 Proteomic analysis of samples

Intracellular and extracellular proteins produced by *S. toxytricini* during growth were extracted from the growth medium and pellets separately. For protein extraction from culture broth, the acetone precipitation method was used and from protein isolation from pellets, pellets were processed in lysis buffer at pH 7.0 [7,21]. Isolated proteins were analyzed by 10% SDS PAGE.

## 3 Results and Discussion

### 3.1 Growth of *S. toxytricini*

The growth of *S. toxytricini* in submerged fermentation showed that if inoculum was transferred from Petri plate to shake flask, it produced diffused mycelia and pellets of smaller size while the transfer of inoculum from shake flask to shake flask, produced visible pellets in the culture broth. Thus it can be assumed that in Petri plate bacterial mycelium was not programmed for pellet formation, but in the culture broth, *S. toxytricini* produces some chemical compound (intracellular or extracellular) that directs the formation of pellets at the submerged level. Microscopic examination of *S. toxytricini*, which was grown using inoculum from a Petri plate revealed that mycelial aggregation took place, but the majority of these aggregations were clumps, loose mycelia, and smaller pellets (upto 20μm). While shake flask to shake flask inoculation produced pellets of larger size (30μm to 2mm) as major morphological form and very less loose mycelia. Thus, it may be assumed that the transformation of mycelia into pellets is influenced by culture conditions, and when *S. toxytricini* is grown in submerged conditions, it produced some bio-chemicals or proteins which controlled pellet formation. Including this heterogeneity of mycelial forms (loose, clump and pellet) is maintained in broth medium which is according to previously reported study (Kumar and Dubey, 2017). Similarly, some studies have reported that variation in culture conditions directly affected pellet morphology of bacterial strain but such mycelial morphology is unwanted for industrial processes (Flardh et al., 2012; Rioseras et al., 2014; van Veluw et al., 2012).

### 3.2 Morphology of pellets

For better insight into pellet structure, the morphology of pellets was analyzed by scanning electron microscope (Fig 1). In this study, it was observed that pellet formation takes place by interwoven hyphae (Fig 1A) where hyphae are joined with each other and form a compact network (Kim and Kim, 2004; Koebsch et al., 2009). The surface of pellet has loose mycelia which are sites of growth and these mycelia undergo fragmentation during agitation to form new pellet (Fig 1B) (Koebsch et al., 2009). Further magnification (2μm scale) revealed that hyphae are attached to each other and form tight junctions (Fig 1C). Microscopic analysis revealed that growing hyphae bears bud-like structures behind the apical tip and these buds like structures form branches which were developed at regular intervals (Fig 1D) (Hempel et al., 2012). Thus it may be assumed that branching behind apical tip and tight junctions assist in compact pellet formation. As reported earlier such types of association protect hyphae from fission and produce multinucleoid structure (Jakimowicz and van Wezel, 2012). The microscopic analysis strongly supported the observations of previous researchers such as; generation of multiple polarity centers (Holmes et al., 2013), growth and development of pellet by tip extension, branching and cross-wall formation (Hempel et al., 2012; Flardh et al., 2012).

**Fig. 1:**
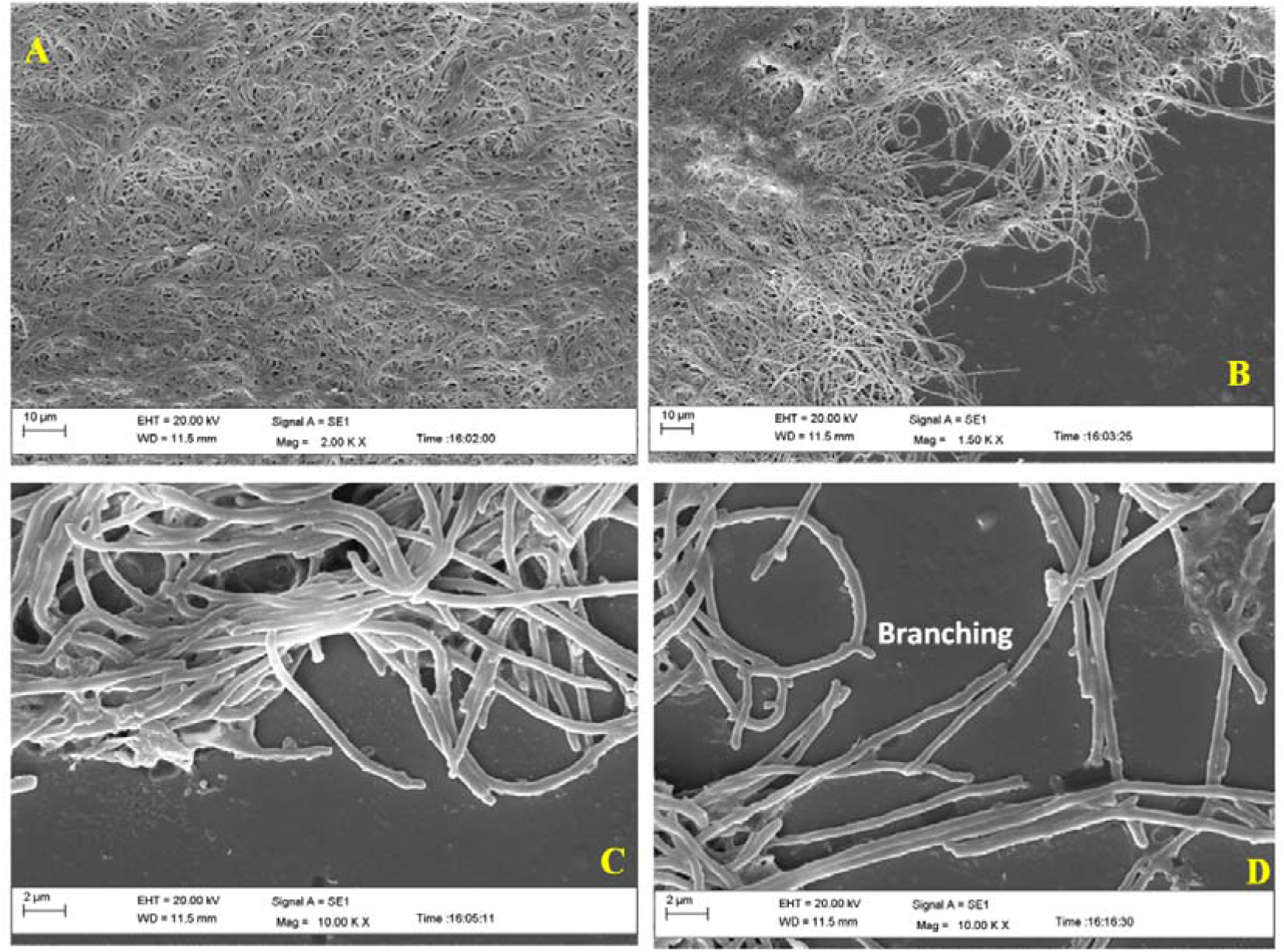
Analysis of pellet morphology and mycelial structure by scanning electron microscopy. **A**-Morphology of pellet at 10μM scale, **B-** Surface of pellet showing free mycelia which show growth and branching at 10μM scale, **C**-Structure of hyphae showing association with each other and branching at growing tips (2μM scale), **D**-Apical growth and branching of hyphae (2μM scale).

### 3.3 Proteomics analysis

Researchers have reported many proteins (DivIVA, Scy, FilP, SsgA, ParA, Tat, CslA, Afsk, CRP, HyaS, TrpM, and Mat complex etc) involved with apical growth, formation of septa, branching in growing hyphae (Ditkowski et al., 2013; Hempel et al., 2012; Holmes et al., 2013; Noens et al., 2007; van Dissel et al., 2018; Vassallo et al., 2020; Willemse et al., 2012). To further elucidate the involvement of different proteins behind pellet formation of *S. toxytricini*, secreted proteins in culture medium and intracellular proteins were analyzed by electrophoresis. In this study, the partial analysis of proteins (intracellular and extracellular) was conducted and It was observed that large numbers of proteins of different molecular weight were produced at intracellular level and extracellular level ranging from 95Kda to less than 20kDa (Fig 2). It is assumed that many of these proteins are involved in pellet formation. It was observed in Fig 1 D, that growing hyphae forms branch behind apical tip and studies have reported the expression of DivIVA (21.5kDA) at growing tip which is suggested to interact with Scy, ParA and FilP to regulate pellet formation (Ditkowski et al., 2013; Flardh et al., 2013; Holmes et al., 2013). It has been suggested that Scy regulate number of polarity centers by associating with DivIVA for new tip construction during branching (Holmes et al., 2013). Including this, Scy was found to sequester DivIVA and initiate the formation of new polarity centers (Holmes et al., 2013). The protein Scy is also assumed to associate with ParA to hyphal tips and control polymerization of ParA. The Scy-ParA association is assumed to be involved in the transition of hyphal elongation into sporulation (Ditkowski et al., 2013). Thus these proteins need to be further analyzed for their involvement in pellet formation. Including this, the presence of other proteins involved in the growth and branching of hyphae into pellet formation need to be analyzed before making any final conclusion.

**Fig. 2:**
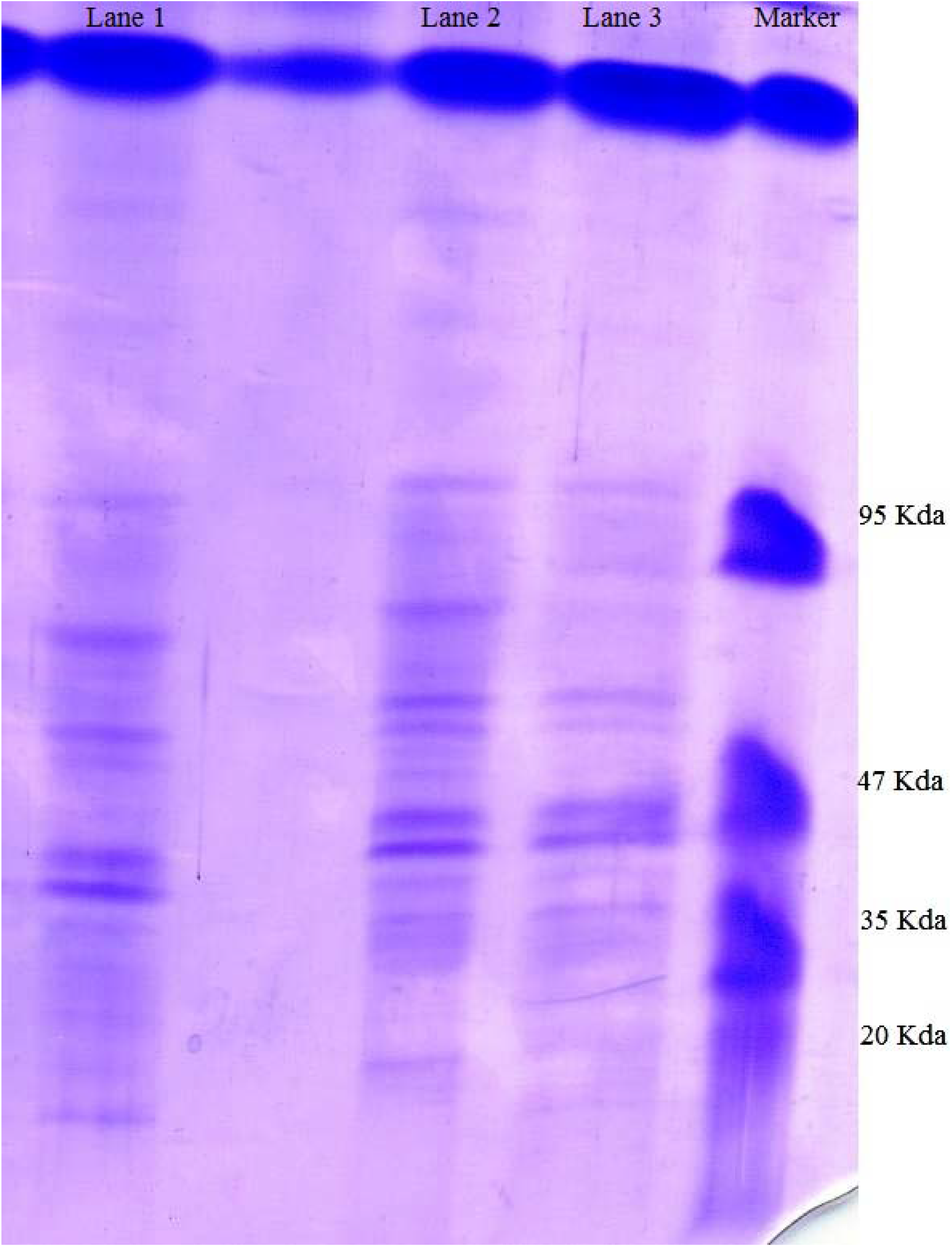
Analysis of extracellular and intracellular proteins of *S. toxytricini* by SDS-PAGE. Lane 1 represents total intracellular proteins, lane 2,3 represents total extracellular proteins isolated from culture medium. Marker lane is molecular weight marker.

### 3.4 Mechanism of pellet formation

In the previous study authors have discussed the process of pellet formation (Kumar and Dubey, 2017) (Fig 3). DivIVA is found associated with cell wall synthesis, genetic competence, and chromosome segregation during sporulation (Labajova et al., 2021). It had been also suggested that DivIVA with some other proteins (Scy, ParA and FilP) regulates formation of polarisome and forms branches in hyphae. Though continuous agitation of culture medium enables clumping of mycelia, but there may be some external biopolymer assisting the association of hyphae with each other resulting compact pellet. To analyze this process, microscopic observation was conducted to understand the pattern of pellet formation. It was revealed that during growth, hyphae perform growth and branching behind apical tip. The growing hyphae come in close proximity to form a network of hyphae which forms a reaction centre like structure. This structure further grows and associates with other hyphae and finally converts into a compact structure (Fig 3). This hypothesis is also supported by the study that apical growth regulates cell polarity which determines the morphology of pellet (Hempel et al., 2012; Holmes et al., 2013). Including this, researchers have demonstrated the formation of cross-walls during growth of streptomycetes which compact the structure into pellet (Flardh et al., 2012). Thus it may be suggested that pellet formation involves the process of hyphae growth, development of polarity, branching and formation of cross-walls which results into transformation of hyphae into compact pellet.

**Fig. 3:**
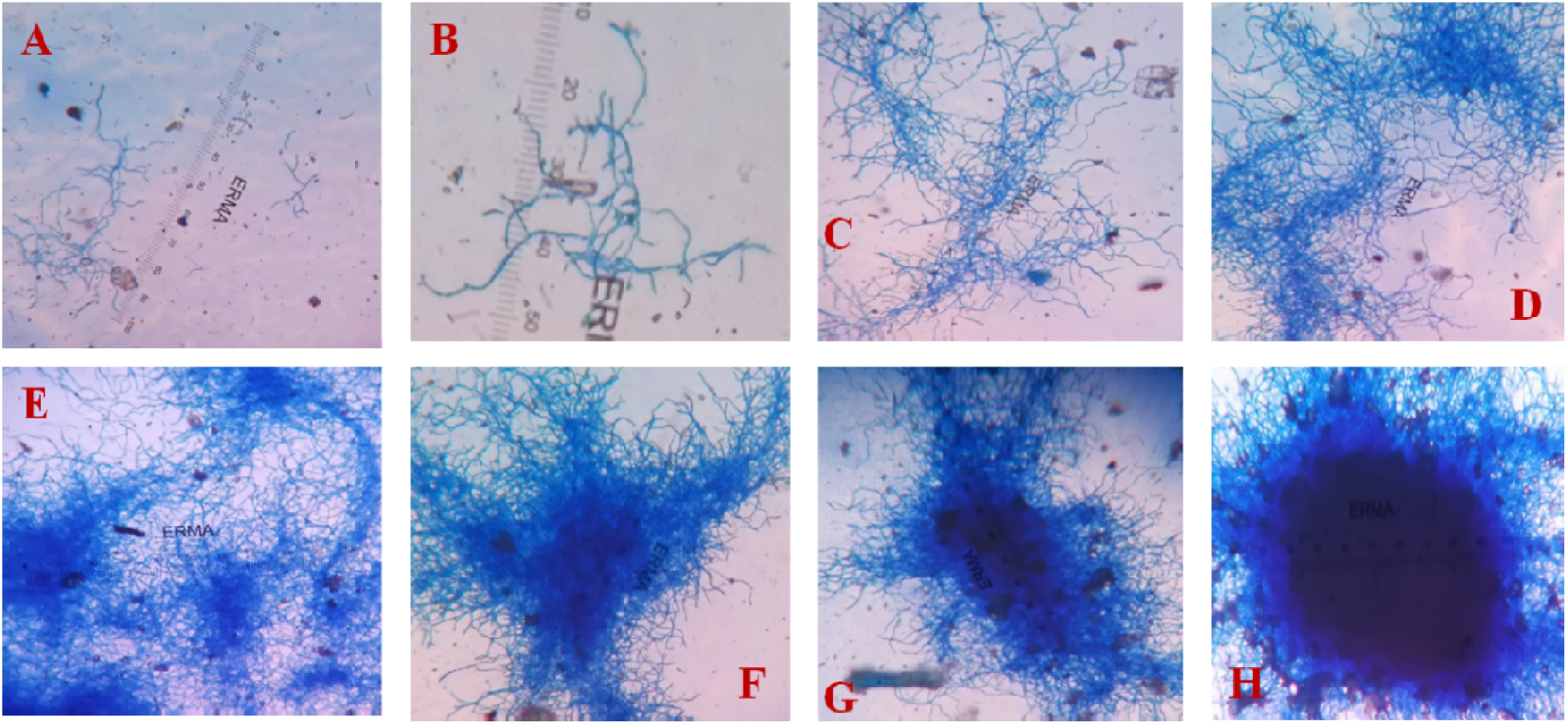
Proposed mechanism for pellet formation. **A**-Diffused mycelia, **B**-Mycelia are coming together, **C**-mycelia are clumping, **D**-mycelial clumps are more visible and a reaction centre type structure is formed which possibly directs pellet formation, **E**,**F**,**G**-mycalial clumps are more condensed, **H**-Pellet formed and periphery having mycelia

## 4 Conclusions

Streptomycetes are filamentous microorganisms which are reported for the production of many valuable natural products including antibiotics and enzymes. The hyphae of these microbes represent a distinct morphological form i.e. pellet during submerged cultivation and undesirable for industrial process. Though, studies have reported production of antibiotics using pellet morphology. Investigators have reported the involvement of culture conditions and biomolecules controlling the morphology of hyphae. In this study, microscopic analysis of morphological forms of *S. toxytricini* at different time revealed that during submerged growth, the growth of hyphae took place by tip extension, branching behind apical tip. The growing hyphae possibly attached with each other and form cross-walls which further transformed into compact pellet (Flardh et al., 2012; Hempel et al., 2012). The partial proteomic analysis in order to understand the role of proteins in pellet formation revealed that there is presence of low weigh proteins in pellet and culture medium which are possibly involved in pellet formation but detailed study of such proteins like DivIVA, Scy, FilP, SsgA, ParA, Tat, CslA, Afsk, CRP, HyaS, and Mat complex etc. is required.

## Acknowledgements

Authors sincerely acknowledge the United States Department of Agriculture for providing *S. toxytricini* NRRL B-5426 and Advanced Instrumentation Research Facility, Jawaharlal Nehru University, New Delhi (India) for scanning electron microscopy of pellets.

